# MultiGML: Multimodal Graph Machine Learning for Prediction of Adverse Drug Events

**DOI:** 10.1101/2022.12.16.520738

**Authors:** Sophia Krix, Lauren Nicole DeLong, Sumit Madan, Daniel Domingo-Fernández, Ashar Ahmad, Sheraz Gul, Andrea Zaliani, Holger Fröhlich

## Abstract

Adverse drug events constitute a major challenge for the success of clinical trials. Several computational strategies have been suggested to estimate the risk of adverse drug events in preclinical drug development. While these approaches have demonstrated high utility in practice, they are at the same time limited to specific information sources and thus neglect a wealth of information that is uncovered by fusion of different data sources, including biological protein function, gene expression, chemical compound structure, cell-based imaging, etc. In this work we propose an integrative and explainable Graph Machine Learning approach (MultiGML), which fuses knowledge graphs with multiple further data modalities to predict drug related adverse events. MultiGML demonstrates excellent prediction performance compared to alternative algorithms, including various knowledge graph embedding techniques. MultiGML distinguishes itself from alternative techniques by providing in-depth explanations of model predictions, which point towards biological mechanisms associated with predictions of an adverse drug event.

**Motivation:** Adverse drug events are a major risk for failure of late-stage clinical trials. Attempts to prevent adverse drug events in preclinical drug development include experimental procedures for measuring liver-toxicity, cardio-toxicity, etc. Yet these procedures are costly and cannot fully guarantee success in later clinical studies, specifically in situations without a reliable animal model. Computational approaches developed for adverse event prediction have shown to be valuable, but are mostly limited to single data sources. Our approach successfully integrates various data sources on protein functions, gene expression, chemical compound structures and more, into the prediction of adverse events. A main distinguishing characteristic is the explainability of our model predictions which allow further insight into biological mechanisms.

## 1. Introduction

Adverse drug events (ADEs) are defined as an injury resulting from the use of a drug, including harm caused by the drug (adverse drug reactions and overdoses) and harm from the use of the drug (including dose reductions and discontinuations of drug therapy) (Nebeker et al., 2004). Experimental approaches to address potential ADEs (e.g. liver-toxicity) based on animal and tissue models are well established in pharma research. Yet, results obtained in such model systems may not always reflect the situation in humans. Furthermore, reliable model systems do not even exist for all indication areas. Computational approaches reflective of human biology could bridge this gap and provide supportive information regarding potential ADE prediction. Hence, there is a strong in interest in computational strategies: ADE prediction has been tackled via statistical methods based on human genetics (Carss et al., 2022; Duffy et al., 2020; Nguyen et al., 2019), chemical structure-based approaches (Liu et al., 2014; Niu & Zhang, 2017; Pauwels et al., 2011; Yamanishi et al., 2012; Zhang et al., 2015; Zhao et al., 2018), approaches using high-throughput literature mining (Deftereos et al., 2011), gene expression data (Cakir et al., 2021; Z. Wang et al., 2016), protein sequences (Takarabe et al., 2012) and electronic health records (Vilar et al., 2012) and data from electronic pharmacovigilance systems such as the FDA Adverse Event Reporting System (Schotland et al., 2021). While each of these approaches have their own merits, they also come along with unavoidable limitations: For example, genetic variants associated with a certain phenotype may not be identifiable in genome-wide association studies due to lack of statistical power. Chemical compound structure can inform about binding affinity to a given target, but does not cover the question whether the choice of a specific target should per se raise safety concerns due to the expected biological downstream consequences. Electronic health records can inform about real-world post-marketing aspects of drugs, but have limited utility in the preclinical drug development phase due to the lack of quantitative biological data.

An alternative strategy is to use biological networks, which represent a rich resource of relational information. In this context knowledge graphs (KGs) have become popular due to their ability to accurately represent multiple types of relationships between different entities (Barabási & Oltvai, 2004; Rebele et al., 2016; Vrandečić & Krötzsch, 2014). That means KGs are multi-relational graphs with entities as nodes and their relations as edges. Relations are represented as triples of (source entity, relation type, target entity). KGs often incorporate a variety of heterogeneous information in the form of different node and edge types. In recent years, numerous knowledge graphs have been published such as OpenBioLink (Breit et al., 2020), Hetionet (D. S. Himmelstein et al., 2017), PharmKG (Zheng et al., 2020) or CTKG (Chen et al., 2021), a knowledge graph on clinical trials. These comprehensive KGs contain a variety of entity types and relation types which model biology as accurately as possible and can be applied to multiple tasks due to their versatile design.

From a network-perspective, ADE prediction can be formulated as a link prediction task in a KG. Earlier approaches extracted manually crafted features of the topology of the KG by using the neighborhood information of each node (Lin et al., 2013) or by extracting local information indexes and path information (Luo et al., 2014). Other authors used an enrichment test of known causes of ADEs to construct features that were subsequently employed in a machine learning algorithm (Bean et al., 2017). Another network-based approach used structural information of the drug molecules for a logistic regression model (Cami et al., 2011). As interactions between co-prescribed drugs are a major cause of ADEs (Aronson, 2015), predicting these by using similarity measures (Fokoue et al., 2016) or by using an approach which generates representative KG embeddings with a convolutional Long short-term memory (LSTM) network (Karim et al., 2019) have been proposed. Another embedding-based approach using node2vec along with a deep neural network was recently developed for predicting ADEs (Joshi et al., 2022). In order to model possible inaccuracies in the KG, Dasgupta *et al*. used confidence scores from natural language processing (NLP) inference for clinical concepts and for edges in the KG during representation learning (Dasgupta et al., 2021).

From a methodological point of view, link prediction in KGs can be addressed by first learning an embedding of the graph structure in Euclidean space. Essentially, the KG embedding is a low-dimensional representation which captures key information about entities and their relations. Typically, entities with similar embeddings are also similar in the original space. Hence, we can assess the likelihood that two entities should be connected by a relation type. In addition to established network representation learning methods such as TransE (Bordes et al., 2013), ComplEx (Trouillon et al., 2016), DistMult (Yang et al., 2014), RotatE (Sun et al., 2019), DeepWalk (Perozzi et al., 2014) and node2vec (Grover & Leskovec, 2016), graph neural networks (GNNs) have emerged as an efficient machine learning method. GNNs were first introduced by Scarselli et al. (2009). Subsequently, graph convolutional neural networks (GCNs) (Kipf & Welling, 2017) and graph attention networks (GATs) (Veličković et al., 2017) were developed as variants of GNNs. GNNs have been successfully applied to various tasks in network analytics, including clustering (Wang et al., 2017) and disease classification (Rhee et al., 2018), prediction of molecular fingerprints (Duvenaud et al., 2015) and protein interfaces (Fout, 2016), as well as prediction of drug-protein interactions (Wu et al., 2022) and poly-pharmacological side effects (Zitnik et al., 2018). A GNN has also been used on a drug-disease graph for ADE prediction (Kwak et al., 2020). This approach has been further developed by combining two GNNs for graph and node embedding in a hybrid approach to predict ADEs via a matrix completion process (Yu et al., 2022). Finally, a graph convolutional autoencoder approach coupled with an attention mechanism has been suggested, leveraging the pairwise attributes for drug-related ADE prediction in a heterogeneous graph (Xuan et al., 2022).

A limitation of these existing KG focused approaches is that they neglect any orthogonal information, including genetic associations, chemical compound structure, gene expression signatures and cell morphology changes. The aim of this paper is thus to address limitations of previous work by developing a Graph Machine Learning approach, which integrates biological networks, genetic variant to phenotype association, gene expression, cell based imaging, protein sequence information, clinical concept embeddings as well as chemical compound fingerprints into one end-to-end trainable algorithm. The idea is thus to integrate a large number of potentially relevant sources of evidence to predict potential ADEs that could occur during clinical trials and therefore reduce the risk of late and costly failures (Schuster et al., 2005). To do so, we built a dedicated KG and designed a novel GNN architecture tailored for ADE prediction. The KG consists of multi-relational and heterogeneous information collected from 14 different databases and is enriched with multi-modal features for each node in order to capture various relevant biomedical data in addition to graph topology. As opposed to state-of-the-art approaches, our proposed MultiGML model is thus designed to integrate multi-modal and in particular also quantitative input data (e.g. gene expression). We demonstrate the superior prediction performance of our approach by comparing it with several state-of-the-art models. Moreover, we introduce a technique to make model predictions explainable, which is crucial in the context of an application in the early phases of drug development. Based on a number of examples we show that our method in this way allows for pointing towards the biological mechanisms associated with a given ADE prediction. Finally, we provide literature evidence for some of the predictions made by our GNN method. The source code and the Python package of MultiGML is available on GitHub (https://github.com/SCAI-BIO/MultiGML).

## 2. Results

### 2.1. Link Prediction Results

In the following, we show the prediction performance of our MultiGML models compared to various competing methods for link prediction in KGs. We first evaluated MultiGML for the task of predicting any link in the KG and second for the more specific task of adverse drug event prediction. Regarding the first task, our MultiGML-RGCN model reached a performance of 0.808 area under precision recall curve (AUPR), and the MultiGML-RGAT model reached an AUPR of 0.798, both outperforming all competing methods (Table 1) by at least ~5%. At the same time precision@1000 was about 0.9 for both variants. When omitting the multi-modal feature embedding (i.e. only learning from the graph topology) the Precision@k of both MultiGML models dropped by 1-2%.

**Table 1.** Model performance results for general relation prediction. The table shows the test results of several competing KG embedding methods, including TransE, RotatE, ComplEx, DistMult and DeepWalk, as well as our two tested MultiGML model variants. Best results are marked in bold. Both RGCN and RGAT variants of the MultiGML model were tested with two types of input features. Basic features are vectors initialized with the Xavier-Glorot method and are unimodal features. Multimodal features refer to the representative features of several modalities for each node type described in section 3.1.2.

| Model | AUROC | AUPR | Precision@1000 |
| --- | --- | --- | --- |
| TransE | 0.667 | 0.633 | 0.715 |
| RotatE | 0.793 | 0.759 | 0.909 |
| ComplEx | 0.757 | 0.699 | 0.622 |
| DistMult | 0.765 | 0.696 | 0.656 |
| DeepWalk | 0.648 | 0.622 | 0.773 |
| MultiGML-RGCN (basic) | 0.847 | 0.787 | 0.881 |
| MultiGML-RGAT (basic) | 0.843 | 0.793 | 0.902 |
| MultiGML-RGCN (multimodal) | **0.859** | **0.808** | 0.901 |
| MultiGML-RGAT (multimodal) | 0.845 | 0.798 | **0.916** |

We focused subsequently on the task of predicting links between compounds and ADEs. For that purpose we employed a version of our MultiGML model for which we specifically optimized hyperparameters on the validation set with respect to the loss for this specific relation type. Once again, all MultiGML variants performed better than all competing methods with AUROC, AUPR and precision@30 all close to 1 (Table 2). Compared to the Random Forest approach by Wang et al. performance gains were highly significant. Notably reported performance measures were based on the negative sampling scheme explained in section 3.3.1. When increasing the ratio of negative samples in the test set from 1:1 to 1000:1 AUROC and AUPR remained stable, and precision@30 was stable for ratios from 1:1 to 100:1 (see Suppl. Figure 1).

**Table 2.** Model performance results for predicting a novel relation between a drug and an ADE. Test results of competing KG embedding methods, including TransE, RotatE, ComplEx, DistMult, DeepWalk, and additionally Random Forest for adverse drug event prediction in comparison to our MultiGML models. Best results are marked in bold. Both MultiGML-RGCN and -RGAT variants were tested with basic and multimodal input features. Basic features are vectors initialized with the Xavier-Glorot method and are unimodal features. Multimodal features refer to the representative features of several modalities for each node type described in section 3.1.2.

| Model | AUROC | AUPR | Precision@30 |
| --- | --- | --- | --- |
| TransE | 0.293 | 0.389 | 0.233 |
| RotatE | 0.943 | 0.915 | 0.967 |
| ComplEx | 0.884 | 0.934 | **1.0** |
| DistMult | 0.963 | 0.966 | **1.0** |
| DeepWalk | 0.575 | 0.604 | 0.567 |
| Random Forest | 0.512 | 0.164 | 0.464 |
| MultiGML-RGCN (basic) | **1.0** | **1.0** | **1.0** |
| MultiGML-RGAT (basic) | **1.0** | **1.0** | **1.0** |
| MultiGML-RGCN (multimodal) | **1.0** | **1.0** | **1.0** |
| MultiGML-RGAT (multimodal) | 0.980 | 0.982 | **1.0** |

We next evaluated the performance for predicting links between genes and phenotypes, which would be of relevance in the context of target selection. For this purpose we used our MultiGML models which were optimized for general link prediction. Once again all variants of the MultiGML model outperformed competing methods with AUROC ~0.89, AUPR ~0.83 and precision@100 ~0.84 (Table 3).

**Table 3.** Model performance results for predicting a novel gene - phenotype association. Test results of competing KG embedding methods, including TransE, RotatE, ComplEx, DistMult and DeepWalk for prediction of a gene - phenotype association in comparison to our MultiGML models. Both MultiGML-RGCN and -RGAT variants were tested with basic and multimodal input features. Basic features are vectors initialized with the Xavier-Glorot method and are unimodal features. Multimodal features refer to the representative features of several modalities for each node type described in section 3.1.2.

| Model | AUROC | AUPR | Precision@100 |
| --- | --- | --- | --- |
| TransE | 0.735 | 0.674 | 0.533 |
| RotatE | 0.723 | 0.680 | 0.710 |
| ComplEx | 0.843 | 0.770 | 0.790 |
| DistMult | 0.848 | 0.767 | 0.776 |
| DeepWalk | 0.654 | 0.630 | 0.761 |
| **MultiGML-RGCN (basic)** | **0.898** | **0.832** | **0.850** |
| MultiGML-RGAT (basic) | 0.897 | 0.831 | 0.841 |
| MultiGML-RGCN (multimodal) | 0.897 | **0.832** | 0.846 |
| MultiGML-RGAT (multimodal) | 0.892 | 0.826 | 0.827 |

We explored which feature modalities contributed most to our model’s predictions. Notably, in both MultiGML variants, available protein and drug features played an important role, i.e. were selected during the hyperparameter optimization (see Suppl. Figure 2). More specifically, the molecular fingerprint of the drugs as well as the gene ontology fingerprint of the proteins were found to be the best choices of node features for the prediction of ADEs with our MultiGML models. Additionally, gene expression signatures of drugs were identified as relevant by the MultiGML-RGCN model.

### 2.2. Use cases

To illustrate the practical use of our MultiGML method we further explored two newly predicted links between drugs and ADEs that were not part of the KG. Furthermore, we show an example of a newly predicted gene - phenotype association. All links have been predicted with probability >70% by MultiGML-RGAT.

#### 2.2.1. Acute Liver Failure As A Predicted Adverse Drug Event of Alendronic Acid

MultiGML predicted a link between alendronic acid (DRUGBANK:DB00630), a bisphosphonate, and acute liver failure (UMLS:C0162557). Alendronic acid is used to prevent and treat osteoporosis (Wishart et al., 2018), and was found to cause liver damage in a patient that was in treatment for osteoporosis (Halabe et al., 2000). Alendronic acid is also listed in the NIH LiverTox lexicon as a “rare cause of clinically apparent liver injury” (Hoofnagle et al., 2013).

We extracted the attention weights for all relations involving acute liver failure and alendronic acid of the MultiGML-RGAT model of the last graph attention layer (see Figure 1 A). Several relations between alendronic acid and proteins, including two tyrosine phosphatases, PTPRS and PTPN4, and the phenotype Paget’s Disease (UMLS:C0029401), were weighted higher than all other direct relations by the MultiGML-RGAT model. Indeed, alendronic acid is used to treat Paget’s Disease of bone, also known as Osteitis Deformans (Reid & Siris, 1999) by inhibiting tyrosine-protein phosphatases (Wishart et al., 2018). Protein tyrosine phosphatase receptor type S (PTPRS) acts as a metastatic suppressor in hepatocellular carcinoma (Z.-C. Wang et al., 2015) and was found dysregulated in cirrhotic liver (Chan et al., 2016). PTPN4 belongs to the same family of proteins and has accordingly been reported as a prognostic marker for hepatocellular carcinoma (Zhangyuan et al., 2018).

**Figure 1.**
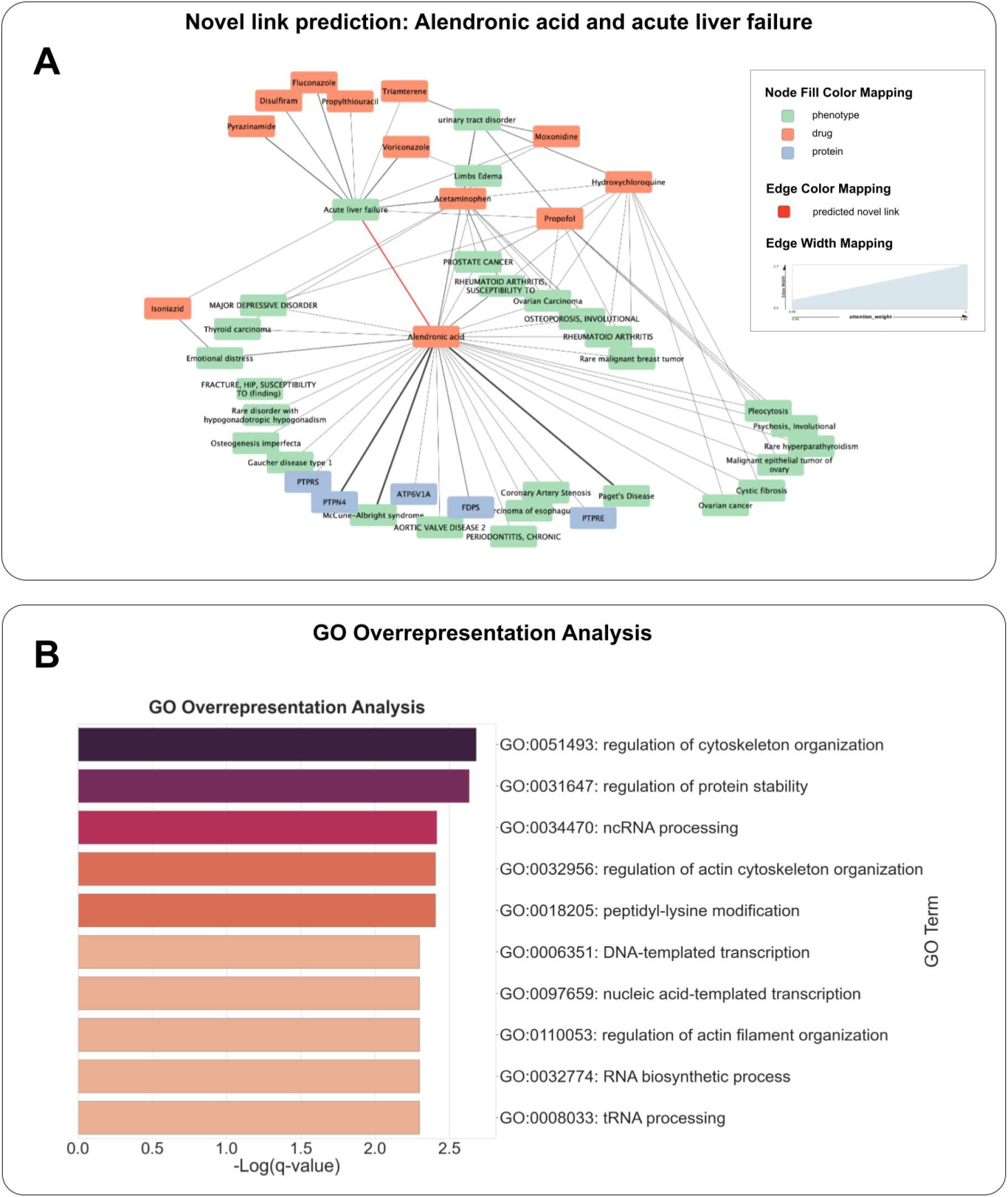
Prediction of acute liver failure as an ADE for alendronic acid. **A)** Novel prediction of acute liver failure (UMLS:C0162557) as a potential ADE of alendronic acid (DRUGBANK:DB00630) in the KG, colored in red. The attention weight for every edge from the last MultiGML-RGAT graph attention layer is indicated by the edge strength. **B)** GO overrepresentation analysis of the top 100 genes from the L1000 drug signature identified via the integrated gradients method. Top 10 enriched terms (one per cluster) created with Metascape. -Log_10_(q) - values are reported and color coded for each term.

To better understand the prediction by our MultiGML model, we investigated the importances of the gene expression signature of the drug alendronic acid in the prediction. More specifically, we identified the top 100 influential genes of the L1000 gene expression signature of alendronic acid according to the integrated gradients methods and performed a Gene Ontology (GO) overrepresentation analysis via a hypergeometric test. After multiple testing correction according to the Benjamini-Hochberg method and choosing a false discovery rate cutoff of 5% we found that regulation of the cytoskeleton organization and protein stability were important for the prediction of acute liver failure as an adverse drug event of alendronic acid (see Figure 1 B, Supp. Table 1). A comprehensive list of the top 100 positively and negatively attributed genes can be found in the Supplementary Material (Suppl. Table 2).

#### 2.2.2. Paralysis As A Predicted Adverse Drug Event of Kanamycin

Kanamycin (DRUGBANK:DB01172), which is an aminoglycoside bactericidal antibiotic (Wishart et al., 2018), was predicted to be associated with a paralytic side effect (UMLS:C0522224) (Figure 2 A iii). Kanamycin is used to treat a wide variety of bacterial infections (Wishart et al., 2018). In several studies, Kanamycin was reported to be neurotoxic and induce neuromuscular paralysis or blockades (Freemon, 1967; Naiman et al., 1965; Pittinger et al., 1970). More recent studies with organoids suggested a damaging effect on early postnatal but not on adult ganglion neurons (Gao et al., 2017). Indeed, ototoxicity of kanamycin is a significant dose-limiting side effect (Heysell et al., 2018).

**Figure 2.**
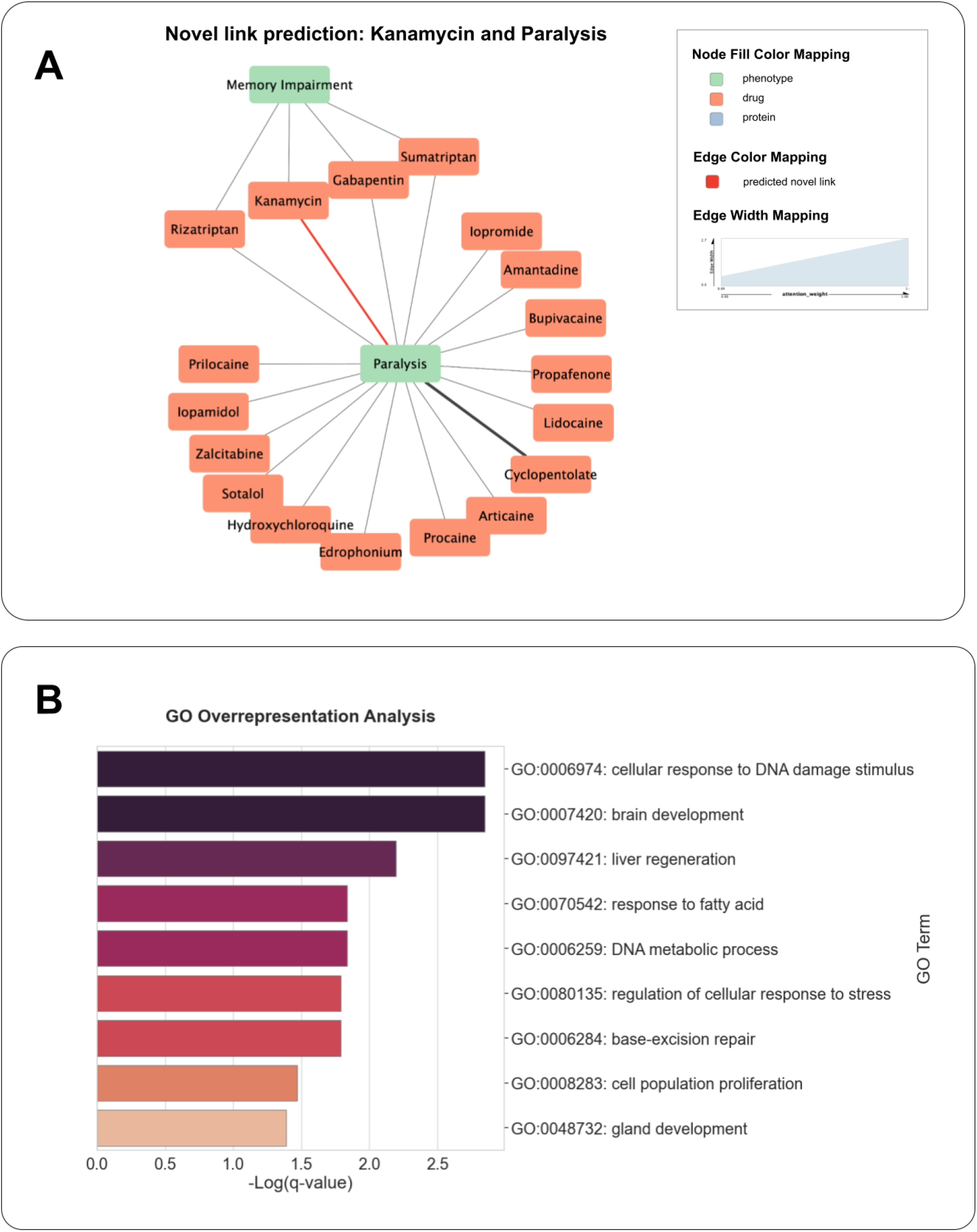
Prediction of paralysis as an ADE for kanamycin. **A)** Novel prediction of paralysis (UMLS:C0522224) as potential ADE of kanamycin (DRUGBANK:DB01172) in the KG, colored in red. The attention weight for every edge from the last MultiGML-RGAT graph attention layer is indicated by the edge strength. **B)** GO overrepresentation analysis of the top 100 genes from the L1000 drug signature identified via the integrated gradients method. Top enriched terms created with Metascape. -Log_10_(q) - values are reported and color coded for each term.

As done previously, we extracted the attention weights for all relations involving kanamycin and paralysis of the MultiGML-RGAT model of the last graph attention layer (see Figure 2 A). Of all direct relations, the relation between the drug cyclopentolate and the phenotype paralysis was weighted higher than all other direct relations by the MultiGML-RGAT model. Cyclopentolate (DRUGBANK:DB00979) is an anticholinergic agent used to dilate the eye for diagnostic and examination purposes (Wishart et al., 2018). Additionally to inducing mydriasis - the dilation of the pupil -, cyclopentolate also causes reversible paralysis of the ciliary muscle by blocking muscarinic receptors (Bartlett & Jaanus, 2008).

We investigated which genes from the gene expression signature of kanamycin were found to be important by the integrated gradient analysis. We performed a GO overrepresentation analysis of the top 100 genes via a hypergeometric test (see Figure 2 B, Suppl. Table 3), reporting the Benjamin-Hochberg adjusted *p*-value (q-value) for multiple testing, and found that biological processes involved in metabolism and responses to stimuli were significantly overrepresented. Altogether, this demonstrates the ability of our method to point towards biological mechanisms associated with ADEs. A comprehensive list of the top 100 genes can be found in the Supplementary Material (Suppl. Table 4).

#### 2.2.3. Association of WNT3 with Thrombophlebitis

*WNT3* (HGNC:12782) is a gene that is part of the Wnt signaling pathway (KEGG:hsa04310) in humans. Wnt proteins are secreted morphogens that are required for basic developmental processes, such as cell-fate specification, progenitor-cell proliferation and the control of asymmetric cell division, in many different species and organs (Kanehisa et al., 2016). Our MultiGML-RGAT model predicted an association between WNT3 and Thrombophlebitis (UMLS:C0040046). Thrombophlebitis is an inflammation of a vein associated with a blood clot (Bodenreider, 2004). Several studies suggest that WNT signaling has a regulatory role in inflammation (George, 2008), is involved in the calcification of vascular smooth muscle cells (Bundy et al., 2021), and that it is a key player in the development of vascular disease, including thrombosis (Foulquier et al., 2018). A recent study on endothelial injury has shown protective effects of Wnt signaling (Y. Wang et al., 2021). More specifically, attenuated apoptosis and exfoliation of vascular endothelial cells and infiltration of inflammatory cells was observed upon activation of the Wnt/beta-catenin pathway.

In Figure 3, we display the novel link of WNT3 to thrombophlebitis in the knowledge graph together with the attention weights learned by our MultiGML-RGAT model. Several relations between WNT3 and other proteins were attributed a high attention weight, including insulin (INS), LRP6, as well as their relations with other proteins, FZD4, FZD7, FZD9, SFRP2 with INS and FZD10 with LRP6. Betamethasone and its relation to thrombophlebitis as an ADE was also attributed with a high attention weight. Associations between the phenotypes essential thrombocythemia (UMLS:C0040028) and rare diabetes mellitus (UMLS:C5681799) with insulin were also much attended by the model. The association between thrombosis, vascular inflammation and diabetes in relation with insulin resistance has been observed in many studies (Pechlivani & Ajjan, 2018; Piazza et al., 2012). A modulation of the interaction of the insulin and Wnt signaling has even been proposed as an attractive target in treating diabetes (Abiola et al., 2009), and could potentially have a role in mediating the effect of inflammatory conditions affecting the vascular system, such as thrombophlebitis.

**Figure 3.**
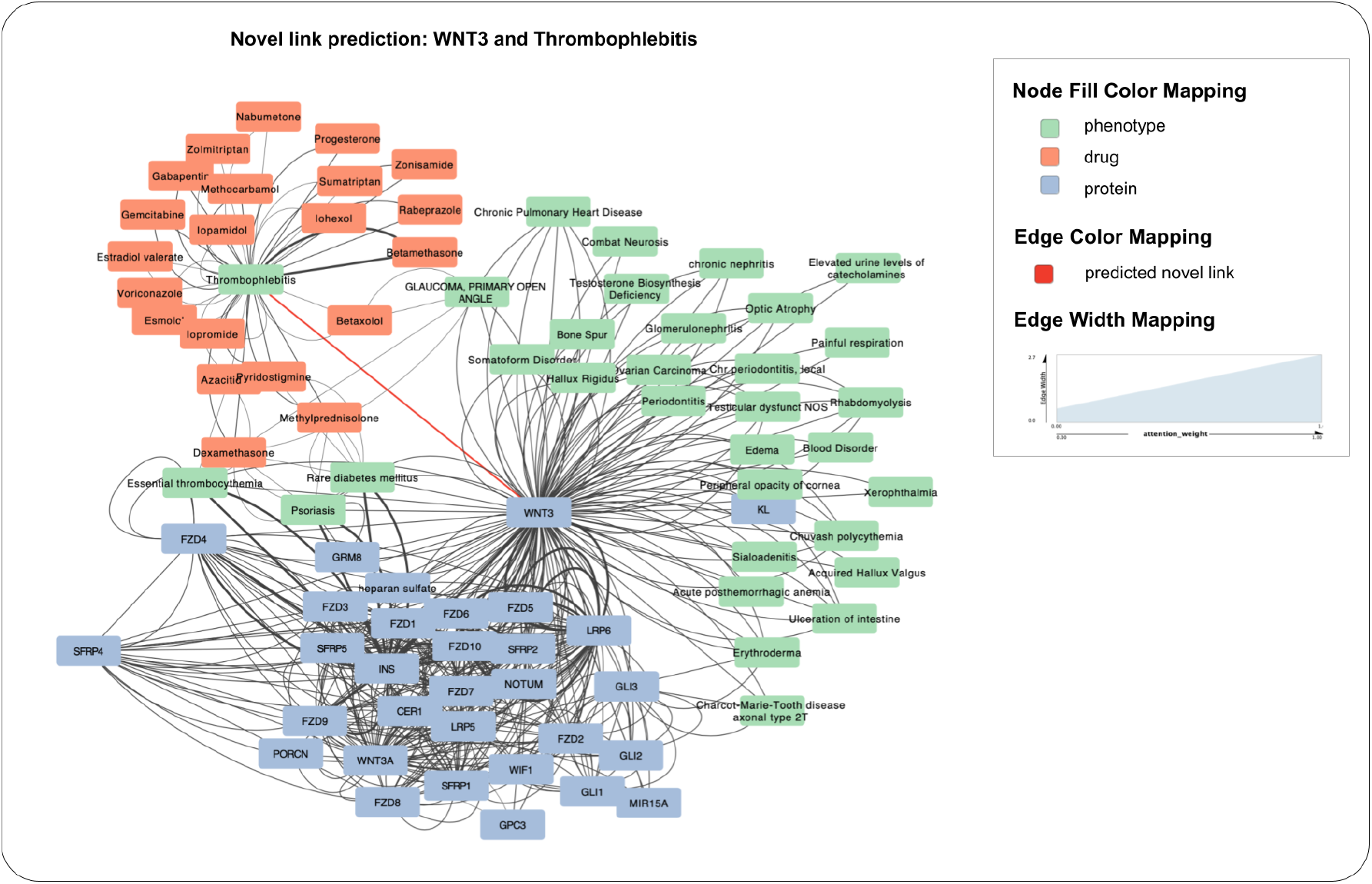
Prediction of thrombophlebitis as a phenotype associated with WNT3. Novel prediction of thrombophlebitis (UMLS:C0040046) associated with WNT3 (HGNC:12782) in the KG, colored in red. The attention weight for every edge from the last MultiGML-RGAT graph attention layer is indicated by the edge strength.

## 3. Materials and Methods

### 3.1. Multi-modal Knowledge Graph Generation

#### 3.1.1. Integration of Biomedical Knowledge from Databases

We integrated information from 14 well-established databases to generate an heterogeneous KG. Our KG contains information about interactions and associations between drugs, proteins, and phenotypes. We introduce the node type ‘phenotype’ to resolve the ambiguity between ADEs and diagnoses. That means ADEs and diagnoses are both subsumed as ‘phenotype’. As a result, we generated a heterogeneous and multi-relational KG with 3 different entity types and 8 different relation types (see Figure 4A). The relation types that occur in the KG are drug-protein, protein-phenotype, genetic variant-phenotype, drug-adverse drug event and 3 different types of protein-protein interactions (physical interaction, functional interaction and signaling interaction) (see Suppl. Table 5). The knowledge graph is formally defined as *G = (N, R*), with *N* as entities, and *R* as relations.

**Figure 4.**
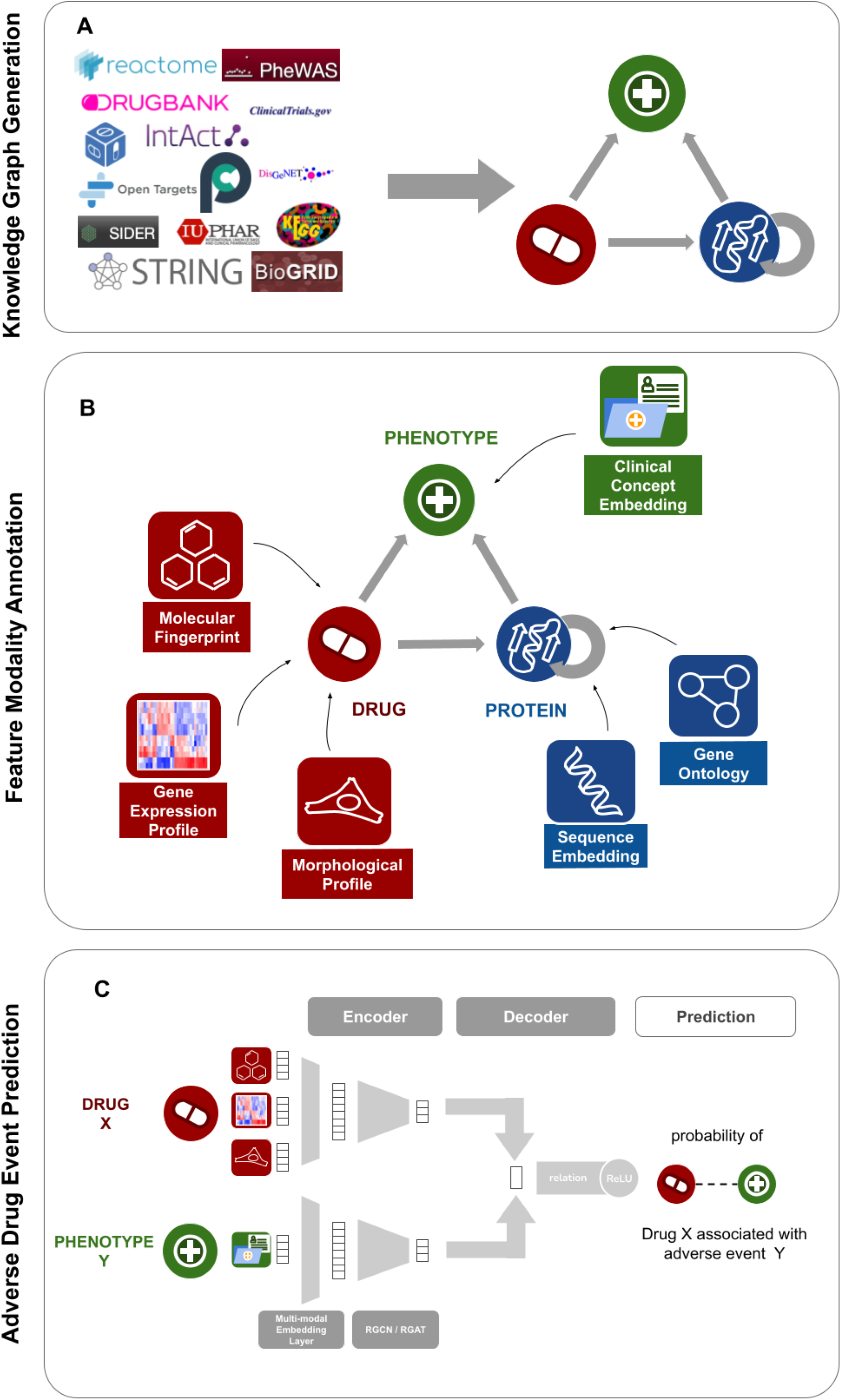
Overview of workflow. A) Knowledge Graph compilation. In the first step of data processing, interaction information from 14 biomedical databases was parsed with data on drug-drug interactions, drug-target interactions, protein-protein interactions, indication, drug-ADE and gene-phenotype associations. The data was harmonized across all databases and a comprehensive, heterogeneous, multi-relational knowledge graph was generated. B) Feature definition. Descriptive data modalities were selected to annotate entities in the knowledge graph. Drugs were annotated with their molecular fingerprint, the gene expression profile they cause, and the morphological profile of cells they induce. Proteins were annotated with their protein sequence embedding and a gene ontology fingerprint. Phenotypes, comprising indications and ADEs, were annotated by their clinical concept embedding. C) Proposed MultiGML approach. The heterogeneous Knowledge Graph with its feature annotations is used as the input for our graph neural network approach, the MultiGML. For each node entity, a multi-modal embedding layer learns a low dimensional representation of entity features. These embeddings are then used as input for either the RGCN or RGAT of the encoder (see section 3.2.1), which learns an embedding for each entity in the KG. A bilinear decoder takes a source and a destination node, drug X and phenotype Y in the example here, and produces a score for the probability of their connection, considering their relation type with each other.

Interaction information between approved drugs and their protein targets was taken from DrugBank (Wishart et al., 2018) and IUPHAR (Harding et al., 2018). Associations between proteins and indications (here: phenotypes) with a high confidence score > 0.6 were extracted from DisGeNet (Piñero González et al., 2020) and specific gene-phenotype associations from PheWAS (Denny et al., 2010), with an odds-ratio > 1. Drug indications for diseases were obtained from OpenTargets (Ochoa et al., 2021) and ClinicalTrials (Zarin et al., 2011). Protein-protein interactions were gathered from renowned databases, including KEGG (Kanehisa et al., 2016; Kanehisa & Goto, 2000), Reactome (Jassal et al., 2020), BioGRID (Oughtred et al., 2021), IntAct (Kerrien et al., 2012), PathwayCommons (Cerami et al., 2011) and STRINGDB (Szklarczyk et al., 2019), where interactions from the latter were filtered for genetic associations with a homology score > 600. The OFFSIDES database (Tatonetti et al., 2012) as well as the renowned SIDER database (Kuhn et al., 2016) were used to extract known ADEs of drugs (see Suppl. Table 6).

Because more severe ADEs tend to increase the risk for a drug to fail in clinical trials or be withdrawn from the market, the following heuristic was employed to filter out more severe ADEs from the information contained in the aforementioned databases: first, we designed a novel metric called the *failure ratio*, which we computed for each phenotype in the graph which served as a target node in an “adverse drug event” edge. The *failure ratio* is defined in Equation 1. Given the number of trials in which a given phenotype is listed as an “adverse drug event” on *ClinicalTrials.gov*, the *failure ratio* is equal to the proportion of these trials which were suspended, terminated, or withdrawn. All phenotypes with a failure ratio greater than 0.75 among at least 3 trials were chosen to be used for the graph. This resulted in nearly two hundred ADE relations, approximately 0.1% of the unique ADE between SIDER and OFFSIDES, which were subsequently added to the KG. All identifiers and interaction types were harmonized across all databases (Figure 4 A). A complete overview about the entity and relation types in the KG is provided in Table 4 and Table 5.

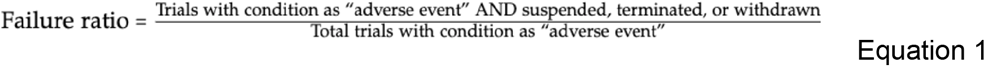

**Table 4.**
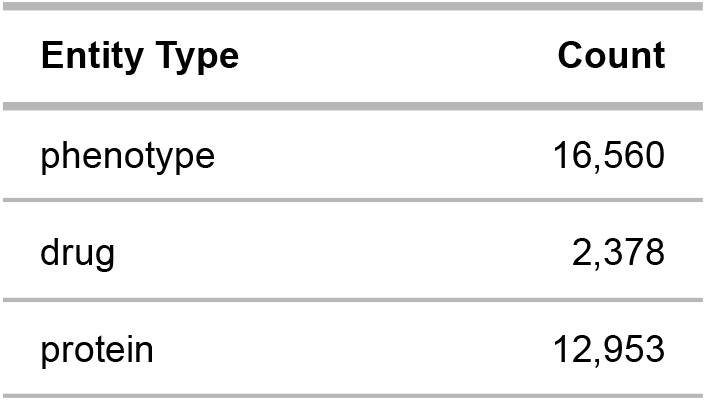
Overview of entities in the knowledge graph.

| Entity Type | Count |
| --- | --- |
| phenotype | 16,560 |
| drug | 2,378 |
| protein | 12,953 |

**Table 5.** Overview of relation types in the knowledge graph.

| Relation Type | Count |
| --- | --- |
| drug-adverse drug event | 197 |
| functional protein-protein association | 1,761 |
| protein-phenotype | 5,448 |
| physical protein-protein interaction | 6,087 |
| drug-protein | 12,451 |
| drug-indication | 12,848 |
| genetic variant-phenotype | 159,202 |
| protein-protein signaling interaction | 222,078 |

#### 3.1.2. Definition of Entity Related Features

Due to the increasing availability of descriptive data modalities for drugs, proteins, and phenotypes, we decided to incorporate multiple feature modalities in our dataset (see Figure 4 B). We chose modalities that were descriptive for the individual entity type, and generated features for each entity type as described below:

- ***DRUGS:*** Transcriptomics data are informative about the effect of a drug on biological processes in a defined system of a cell culture experiment. A molecular signature can therefore be generated for each drug, measuring the gene expression fold change of selected transcripts. We chose the LINCS L1000 dataset (Duan et al., 2014) to annotate the drugs with gene expression profile information. More specifically, we retrieved the consensus signatures calculated by Himmelstein et al. (D. Himmelstein et al., 2016; D. S. Himmelstein et al., 2017). The background is that each LINCS compound may have been assayed across multiple cell lines, dosages and replicates. Himmelstein et al. thus estimated a single consensus transcriptional profile across multiple signatures (Himmelstein, Daniel, 2015) The effects of a drug perturbation in a cell culture experiment can not only be seen in the gene expression fold change, but also in the change in morphology of the treated cells. Therefore, we additionally annotate the drug with the Cell Painting morphological profiling assay information from the LINCS Data Portal (LDG-1192: LDS-1195) (Schreiber, 2014). Furthermore, the molecular structure of the drugs was also taken into account by generating the molecular fingerprints. We here took the Morgan count fingerprint (Rogers & Hahn, 2010) with a radius = 2, generated with the <u>RDKit</u> (Landrum, 2010).
- ***PROTEINS:*** We used structural information of proteins by utilizing the protein sequence embeddings. We generated the embeddings for each protein with the ESM-1b Transformer (Rives et al., 2021), a recently published pre-trained deep learning model for protein sequences. In addition, we generated a binary Gene Ontology (GO) fingerprint for biological processes for each protein using data from the Gene Ontology Resource (Ashburner et al., 2000; Gene Ontology Consortium, 2021). A total of 12,226 human GO terms of Biological Processes were retrieved and their respective parent terms obtained. This resulted in a 1,298 dimensional binary fingerprint for each protein, with each index either set to 1, if the protein was annotated with the respective GO term or 0 if not.
- ***PHENOTYPES:*** Medical concept embeddings from Beam et al. (Beam et al., 2020) were used to annotate phenotypes including ADEs and indications. The so-called *cui2vec* embeddings were generated on the basis of clinical notes, insurance claims, and biomedical full text articles for each clinical concept. Briefly, the authors mapped ICD-9 codes in claims data to UMLS concepts and then counted co-occurence of concept pairs. After decomposing the co-occurrence matrix via singular value decomposition, they used the popular word2vec approach to obtain concept embeddings in the Euclidean space. We refer to Beam et al. for more details.

### 3.2. Graph Neural Network Architecture

MultiGML consists of two main structures, an encoder and a decoder. The encoder has two main components which create a low-dimensional embedding of each node in the KG (see section 3.2.1.). Due to its design, the encoder can handle multimodal input data. The second part of the model decodes edges from node embeddings with a bilinear form (see section 3.2.2.). The entire model architecture is shown in Figure 4C and discussed in more detail in the subsequent paragraphs.

#### 3.2.1. Encoder

##### 3.2.1.1. Multi-modal Embedding of Node Features

Due to the multi-modal character of our KG, we require a model that can integrate input features of multiple modalities for one node into the message passing. To do so, we implemented a specific architecture based on the work by (Lemsara et al., 2020) that combines representations from different data modalities (Figure 5).

**Figure 5.**
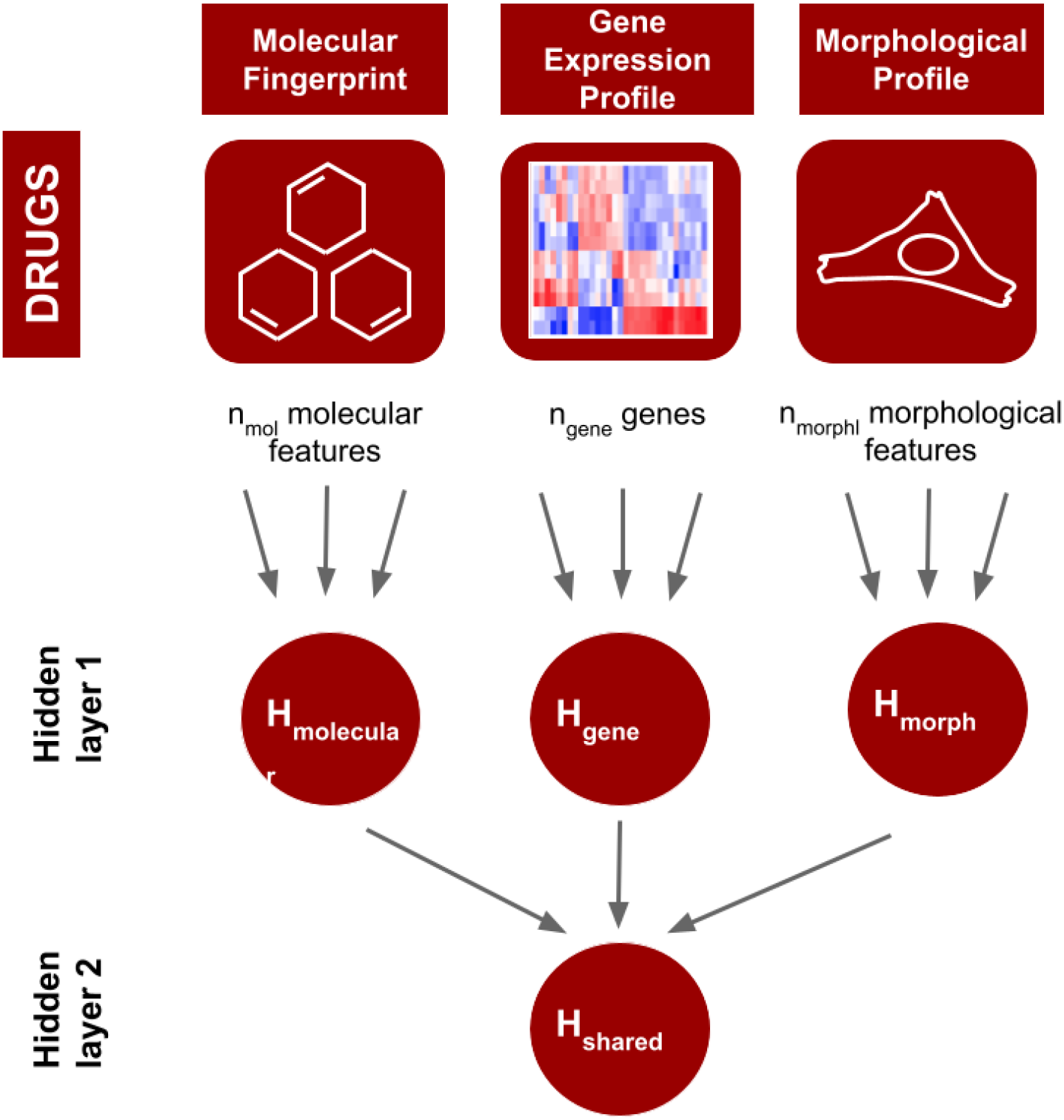
Multi-modal embedding. (example of drug input): Each drug is represented by *f* different feature modalities, which are fed into a multi-modal neural network with bottleneck architecture. That means *H_molecular_, H_gene_, H_morph_*, are the output of dense feed-forward layers, each having *k_r_*/2 hidden units, where *k_r_* is the number of original input features for data modality *r*. *H_shared_* = (*H_molecular_* ||*W_gene_* ||*W_morph_*) represents the multi-modal embedding.

In a nutshell, this multimodal embedding learns hidden representations of each modality separately in the first densely connected layer. The hidden feature representations are then concatenated and passed to a second densely connected layer to generate a shared multimodal embedding for each entity *v* in the KG: Let *x*_1_, *x*_2_,…, denote the *k* feature vectors of dimensions *d*_1_, *d*_2_, …, *d_k_* associated to entity *v*. The embedding *H_shared_* is therefore learned as follows:

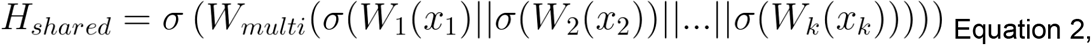

where *σ* is the tanh activation function and || denotes a concatenation.

We use dropout units in each layer with a dropout ratio that is adjusted during Bayesian hyperparameter optimization. This is followed by a batch normalization with a tanh activation function.

##### 3.2.1.2. Relational Graph Convolutional Neural Network (RGCN)

KGs often incorporate a variety of heterogeneous information in the form of different node and edge types. In the following, we will refer to the prominent Relational Graph Convolutional Neural Network (RGCN) that was proposed by Schlichtkrull et al. (2018) to deal with the multirelational data characteristic of KGs. The RGCN includes information from the neighborhood of a node into the message passing by differentiating between the relation types. Due to this characteristic, the model is able to learn the inherent relationships between the entities in the KG.

The RGCN takes as input a heterogeneous multi-relational knowledge graph *G* with features 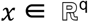 and learns an embedding *h_i_* of each entity *v_i_* ∈ *N* in the KG. The architecture of the implemented model has three consequent RGCN layers (Schlichtkrull et al., 2018)I:

- **input to hidden layer:** input feature vectors 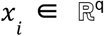 are transformed into their hidden representation 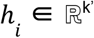
- **hidden to hidden layer**: convolution of hidden feature vectors 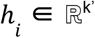, maintaining their shape
- **hidden to output layer**: hidden feature vectors 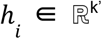 are transformed into their latent representation 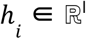

The message passing for node *i* is given by:

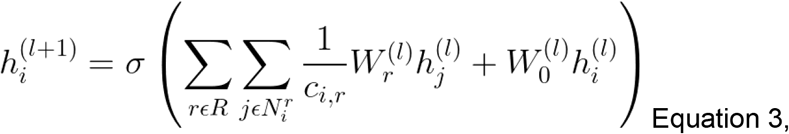

The updated hidden representation *h_i_* of entity *v_i_* at layer *l* + 1 is a non-linear combination of the hidden representations of neighboring entities with index 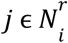. weighted by the learnable relation type specific weight matrix 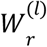. Here, *N_r_*, is the set of neighbors of node *v* of relation type *r*. A self-loop is defined by adding the node’s own hidden representation *h*, multiplied by the weight matrix *W*_0_. *c_i,r_* is a normalization constant that is task-dependent and can either be learned or chosen in advance, such as 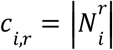. We refer to this variant of our MultiGML model as MultiGML-RGCN.

##### 3.2.1.3. Relational Graph Attention Network (RGAT)

Alternatively to the RGCN we considered a relational graph attention network (Veličković et al., 2017) as part of the encoder. The input is a set of entity features *h* = {*h*_1_, *h*_2_, *h*_3_, …, *h_N_*], *h_i_* ∈ *R^q^* with *N* being the number of entities and *q* being the number of features of each entity. Self-attention is performed on the entities, whereby a shared attention mechanism *a* computes attention coefficients (Veličković et al., 2017) for each relation type (Equation 4)

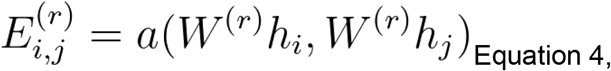

The attention mechanism *a* is a single-layer feedforward neural network feeding into a Leaky ReLU unit (angle of negative slope = 0.2). The attention coefficients are normalized across all choices of *j* via the softmax function (Busbridge et al., 2019; Veličković et al., 2017),

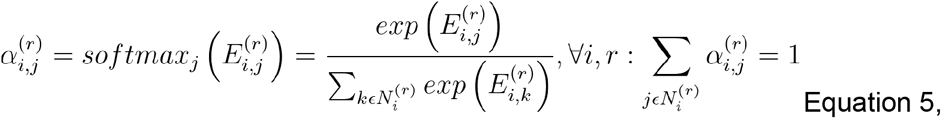

leading to the propagation model for a single node update in a multi-relational graph of the following form (Equation 6):

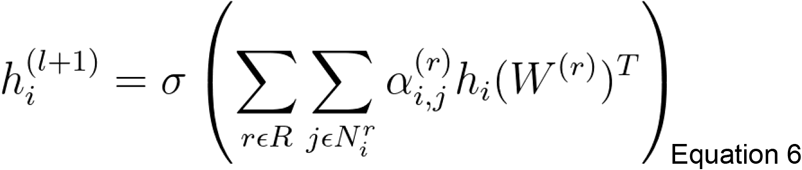

The graph attention layer allows assigning different importances to nodes of a same neighborhood which can be analyzed to interpret the model predictions (Veličković et al., 2017). We refer to this variant of our MultiGML model as MultiGML-RGAT.

#### 3.2.2. Bilinear Decoder

The decoder structure in our model uses the entity embeddings to decode relations in the KG. We calculate a score for a given triple of entities *v_i_* and *v_j_* connected by relation *r*. We use a bilinear form on the embeddings *h_i_* and *h_j_*, with a trainable matrix *M_r_* representing the relation type and apply a sigmoid function σ to the result as in equation 7:

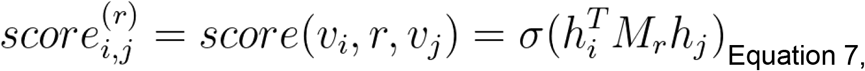

### 3.3. Empirical Evaluation

#### 3.3.1. Model Training Strategy

We trained our MultiGML model on the KG described in section 3.1. The binary cross-entropy loss was applied to supervise the model. We performed a stratified random split of all relations into a 70% train, 20% test and 10% validation set. The stratification was done such that each data subset contained the same fraction of each relation type. The number of relations amounted to 604,903 in the training set, 67,212 in the validation set and 168,029 in the test set. Note that all those real existing relations provide positive examples. An important question in Graph Machine Learning is, how to generate negative samples for non-existing relations. In this work we performed negative sampling where for each positive relation one negative relation was generated by randomly exchanging the target for each source entity according to a uniform distribution (uniform sampling). We employed a hyperparameter optimization with the Optuna package (Akiba et al., 2019), with a customized search space for each hyperparameter (see Suppl. Table 7). Notably, hyperparameter optimization also included the selection of node feature modalities. That means we allowed entire data modalities to be dropped from the model during training. The Tree Parzen Estimator (Bergstra et al., 2011), an independent sampling algorithm, was used for an efficient exploration of the search space. Each hyperparameter optimization for both the RGCN and RGAT model consisted of 50 trials. Models within each trial were trained for 100 epochs unless the hyperband pruner (Li et al., 2016) determined that a trial should be pruned. After each training epoch the model was evaluated on the validation test. After the best set of hyperparameters was found, we trained a final model for 100 epochs with the selected hyperparameters (see Suppl. Table 7), unless early stopping was triggered by a stagnation in the loss calculation for 10 epochs, and then tested the model on the unseen test set.

#### 3.3.2. Comparison against competing methods

We benchmarked MultiGML against several competing approaches for both general link prediction and adverse drug event prediction. All methods were evaluated using the same data splits. We compared our models against four well-established KG embedding approaches, namely TransE (Bordes et al., 2013), RotatE (Sun et al., 2019), DistMult (Yang et al., 2014) and ComplEx (Trouillon et al., 2016) and DeepWalk (Perozzi et al., 2014). Deep Walk is a KG embedding technique based on random walks. DeepWalk creates a mapping of entities to a low-dimensional space of features using a random walk strategy. The entity embeddings and the scores for all samples were generated using the implementation by (Hu et al., 2020), which incorporates a multi-layer perceptron with three layers as a predictor. We also compared our models to a Random Forest (RF) based machine learning approach for ADE prediction (Zhao et al., 2018), which uses gene expression as data as well as compound fingerprints.

### 3.4. Making Models Explainable

From an application perspective it is important to be able to explain which features of a compound influence the model’s predictions in each single instance. For this purpose we build on the integrated gradients method (Sundararajan et al., 2017) as implemented in captum.ai (Kokhlikyan et al., 2020). Integrated gradients is an axiomatic attribution method which represents the integral of gradients with respect to inputs along the path from a given baseline to input (Sundararajan et al., 2017). The integrated gradient along the i^th^ dimension from baseline *x*_0_ to input *x* is defined as

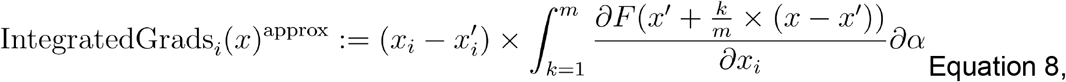

with α as a scaling coefficient, *m* as the number of steps, and *F* as a function 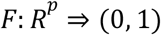 which represents our MultiGML model. For each predicted link between a drug and a side effect, we calculated the integrated gradients to receive the importances of the individual features. We used n = 500 steps for the approximation method. For the baseline *x*_0_, we used a vector of zeros in case of the drug molecular fingerprint. For the drug morphological (i.e. cell painting assay) as well as gene expression features we employed the overall mean vector as baseline. In a further step, the attributions of the Integrated Gradient analysis were evaluated. For the molecular fingerprint of the drugs we decoded the most important molecular moieties from the bits of the molecular fingerprint using RDKit (Landrum, 2010). A gene ontology enrichment analysis of biological processes was performed on the top 100 positively and negatively attributed genes from the molecular gene expression signature of the drugs using Metascape (Zhou et al., 2019). For the enrichment analysis, all genes in the human genome were used as background, and a *Q*-value cut-off of 0.05, a minimum count of 3 and an enrichment factor > 1.5 were chosen. A hypergeometric test was performed and q-values were calculated using the Benjamini-Hochberg procedure (Hochberg & Benjamini, 1990) to account for multiple testing.

### 3.5. Performance Metrics

We evaluate our models on the independent test set according to area under the ROC curve (AUROC), area under the precision-recall curve (AUPR) and precision among the top k predictions (precision@k). Precision@k were calculated based on a ranking of the models’ predicted link probabilities. Due to drastic differences in the number of relation types in the test set, we here report precision@k with k = 1000 for evaluating general link prediction performance, k = 30 for drug - ADE, and k = 100 for gene - phenotype predictions.

## 4. Discussion

We proposed a novel Graph Machine Learning neural network approach for adverse drug event prediction that combines multi-modal quantitative data with a heterogeneous, multi-relational KG. MultiGML demonstrated excellent prediction performance in comparison to a broad set of competing approaches. Furthermore, we demonstrated that predictions made by our MultiGML method can be explained via the method of integrated gradients and visualization of attention weights. We showed that in this way it is possible to identify biologically plausible mechanisms associated with predicted ADEs. Therefore, our approach could provide valuable information during the early phases of drug development, where it is important to lower the failure risk of later clinical trials. Getting insights into relevant biological mechanisms associated with a high risk could support selection of safe targets. We thus see the value of integrating multimodal data into MultiGML not so much in terms of increase in link prediction performance, but much more with regard to the far better interpretability compared to purely graph topology based techniques.

## 5. Limitations

Of course, MultiGML is not without limitations: for example, specific feature modalities may be unavailable for some of the entities in the KG. The neighborhood aggregation approach of MultiGML in such a case provides a way to mitigate this issue, because it essentially smoothens features over the neighborhood of a given entity, but that can not perfectly replace missing information. Furthermore, ADEs are in reality also dependent on pharmacodynamic (PD) properties of a compound, including dose, which are currently not considered in our model. Finally, model explanations can only disentangle model predictions, but they do not always point to the right biological cause of an ADE. Due to the versatile model design, MultiGML offers the perspective of applications of link prediction tasks other than the one discussed in this paper, including drug repositioning. Moreover, MultiGML could be used to integrate other or additional data modalities, for example protein and tissue expression and pathology imaging slides. Altogether, we thus see MultiGML as a flexible approach to support important decisions in early drug discovery.

## Acknowledgements

We thank Bruce Schultz and Aliaksandr Masny for their support during the project. This work was supported by the Research Center Machine Learning (FZML) of the Fraunhofer Cluster of Excellence Cognitive Internet Technologies CCIT.

## Author Contributions

Designed the project: HF, AZ; supervised the work: HF, SM, DDF, AZ, SG; SK and DDF compiled the knowledge graph; SK implemented the models with the help of SM, DDF and AA; SK trained the models with the help of SM; LdL implemented the Random Forest model and SK trained and evaluated the model; SK benchmarked the models; SK, HF, AZ and SG interpreted the results. All authors have read and approved the current version of the manuscript.

