## Supplementary Material for "MultiGML: Multimodal Graph Machine Learning for Prediction of Adverse Drug Events"

#### Supplement

**Supplement 1.** Consensus signature generation.

Pairwise correlations between signatures were calculated and the mean correlation with other signatures was calculated for each signature The similarities were scaled to sum to 1, then the z-score signature vectors were multiplied by their similarity weights. The weighted z-score signature vectors were summed up and used as the consensus signature for DrugBank small molecules (Himmelstein, Daniel, 2015).

#### Figures

**Supplementary Figure 1.** Performance of MultiGML variants for increasing negative samples for side-effect prediction. The performance metrics for AUROC, Average Precision (AUPR) and Precision at k with k=30 on side-effect prediction are shown for both Multi-GML variants, MultiGML-RGCN on the left, and MultiGML-RGAT on the right. For each number of negative samples per positive sample $k \in [1, 5, 10, 50, 100, 250, 500, 750, 1000]$ testing was performed 5 times. The standard deviation is shown in the plots.

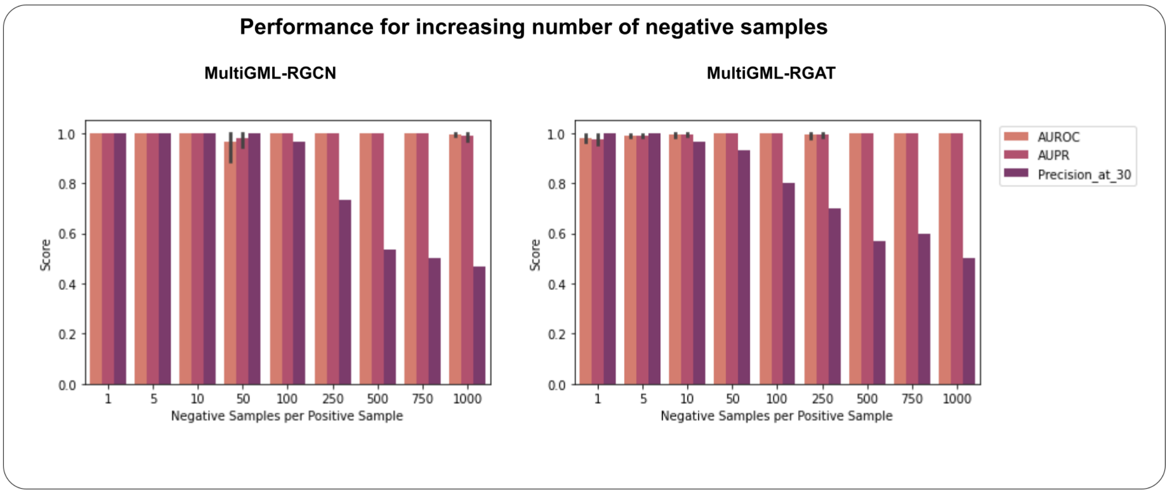

**Supplementary Figure 2.** Hyperparameter importances for parameter during hyperparameter optimization for A) MultiGML-RGCN and B) MultiGML-RGAT. Plots were made with optuna.

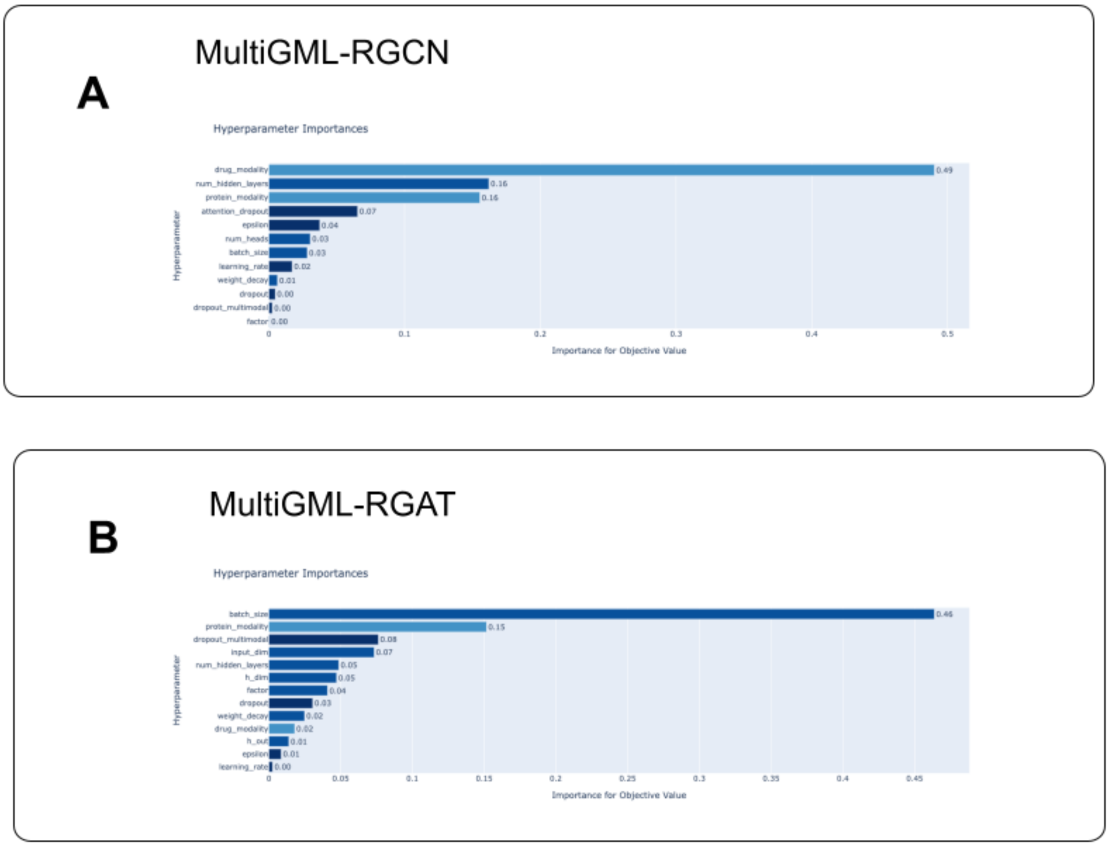


### Tables

**Supplementary Table 1.** GO enrichment analysis of biological processes for the prediction of alendronic acid and acute liver failure. Top 20 clusters with their representative enriched terms (one per cluster) created with Metascape. "Count" is the number of genes in the user-provided lists with membership in the given ontology term. "%" is the percentage of all of the user-provided genes that are found in the given ontology term (only input genes with at least one ontology term annotation are included in the calculation). "Log10(P)" is the p-value in log base 10. "Log10(q)" is the multi-test adjusted p-value in log base 10.

| GO | Description | Count | % | Log10(P) | Log10(q) |
|---|---|---|---|---|---|
| GO:0051493 | regulation of cytoskeleton organization | 17 | 8.50 | -6.87 | -2.68 |
| GO:0031647 | regulation of protein stability | 13 | 6.50 | -6.52 | -2.63 |
| GO:0034470 | ncRNA processing | 14 | 7.00 | -6.13 | -2.42 |
| GO:0018205 | peptidyl-lysine modification | 12 | 6.00 | -5.90 | -2.41 |

| | | | | | |
|---|---|---|---|---|---|
| GO:0006351 | DNA-templated transcription | 16 | 8.00 | -5.59 | -2.30 |
| GO:0009895 | negative regulation of catabolic process | 11 | 5.50 | -4.85 | -1.85 |
| GO:0006325 | chromatin organization | 15 | 7.50 | -4.83 | -1.85 |
| GO:0071229 | cellular response to acid chemical | 6 | 3.00 | -4.54 | -1.61 |
| GO:0030097 | hemopoiesis | 15 | 7.50 | -4.50 | -1.59 |
| GO:0030036 | actin cytoskeleton organization | 13 | 6.50 | -4.12 | -1.32 |
| GO:0006388 | tRNA splicing, via endonucleolytic cleavage and ligation | 3 | 1.50 | -4.11 | -1.32 |
| GO:0022613 | ribonucleoprotein complex biogenesis | 12 | 6.00 | -4.05 | -1.32 |
| GO:0045785 | positive regulation of cell adhesion | 12 | 6.00 | -3.99 | -1.32 |
| GO:0043542 | endothelial cell migration | 5 | 2.50 | -3.98 | -1.32 |
| GO:0043414 | macromolecule methylation | 9 | 4.50 | -3.98 | -1.32 |
| GO:0051603 | proteolysis involved in protein catabolic process | 14 | 7.00 | -3.97 | -1.32 |
| GO:1904874 | positive regulation of telomerase RNA localization to Cajal body | 3 | 1.50 | -3.91 | -1.30 |
| GO:0010810 | regulation of cell-substrate adhesion | 8 | 4.00 | -3.90 | -1.30 |
| GO:0098655 | cation transmembrane transport | 14 | 7.00 | -3.77 | -1.20 |
| GO:0006820 | anion transport | 11 | 5.50 | -3.72 | -1.18 |

**Supplementary Table 2.** Genes of gene expression signature from alendronic acid that were found to be highly positively or negatively attributed and their attribution score. The top 100 negatively and positively attributed genes are listed.

| Entrez Gene ID (positively attributed) | IG attribute score | Entrez Gene ID (negatively attributed) | IG attribute score |
|---|---|---|---|
| GeneID:8270 | 0.0008995434594862807 | GeneID:79991 | -0.00022609830045582771 |
| GeneID:10818 | 0.0006192149080605368 | GeneID:8767 | -0.00022696953517483613 |
| GeneID:8266 | 0.0005135424521145966 | GeneID:8671 | -0.00022726055071385894 |
| GeneID:2115 | 0.0005086604545451612 | GeneID:100133941 | -0.00022778707316336734 |
| GeneID:5613 | 0.0004933134851079761 | GeneID:11062 | -0.00022831795333338567 |
| GeneID:8607 | 0.0004888437663445203 | GeneID:9650 | -0.0002285026585543125 |
| GeneID:323 | 0.000479714563022701 | GeneID:23223 | -0.000230089399928658 |
| GeneID:10165 | 0.0004775488374604353 | GeneID:29100 | -0.00023059142980525314 |
| GeneID:79643 | 0.0004667461276415183 | GeneID:55379 | -0.0002317155058196544 |
| GeneID:3964 | 0.0004540885843581498 | GeneID:8994 | -0.0002322113961224113 |
| GeneID:28972 | 0.0004499720020473656 | GeneID:5501 | -0.00023270193574622522 |

| | | | |
|---|---|---|---|
| GeneID:10362 | 0.0004446570083222257 | GeneID:23370 | -0.0002329637862120419 |
| GeneID:2131 | 0.0004392332159276795 | GeneID:1389 | -0.00023396015149374343 |
| GeneID:23220 | 0.00042694559786747474 | GeneID:81605 | -0.00023475954324415508 |
| GeneID:6881 | 0.0004148178621251951 | GeneID:28962 | -0.00023495287595794428 |
| GeneID:63875 | 0.0004105340282162946 | GeneID:27340 | -0.000235966767954977 |
| GeneID:54512 | 0.0004102499925159137 | GeneID:51582 | -0.0002364244446109688 |
| GeneID:8569 | 0.0003951798959150862 | GeneID:5709 | -0.00023874839244842727 |
| GeneID:9289 | 0.0003851491817799898 | GeneID:25901 | -0.0002395699583677032 |
| GeneID:3182 | 0.00037852541869558406 | GeneID:10360 | -0.0002396031321191957 |
| GeneID:6596 | 0.0003782558882481143 | GeneID:79763 | -0.00023983217670731533 |
| GeneID:3930 | 0.00037757946556041635 | GeneID:65258 | -0.0002404346803024866 |
| GeneID:7422 | 0.00037071445266386924 | GeneID:151011 | -0.0002417044032186267 |
| GeneID:24149 | 0.0003646547605455143 | GeneID:10126 | -0.00024283211230190484 |
| GeneID:2960 | 0.00036237481671281223 | GeneID:54440 | -0.00024337408990114009 |
| GeneID:8192 | 0.00035673952356988137 | GeneID:8148 | -0.00024379506957160647 |
| GeneID:23304 | 0.00035434660466432475 | GeneID:5636 | -0.00024541604181013075 |
| GeneID:55173 | 0.00034364603821231 | GeneID:23325 | -0.0002457759459000078 |
| GeneID:9788 | 0.00034052652177369664 | GeneID:26133 | -0.0002464948043059844 |
| GeneID:55027 | 0.0003397633126819151 | GeneID:5412 | -0.0002473416633911184 |
| GeneID:5870 | 0.00033439205063118114 | GeneID:8558 | -0.00024736601878430983 |
| GeneID:29994 | 0.0003329920458052786 | GeneID:55740 | -0.0002481856756041328 |
| GeneID:10565 | 0.00032835317326553135 | GeneID:8349 | -0.0002482960411639851 |
| GeneID:624 | 0.00032366259164699605 | GeneID:949 | -0.0002490416919672065 |
| GeneID:2316 | 0.00032326800679876157 | GeneID:71 | -0.0002514311204311707 |
| GeneID:28974 | 0.00032310573761873906 | GeneID:3508 | -0.0002533838597956925 |
| GeneID:63933 | 0.00032286788013682053 | GeneID:9829 | -0.00025465704150274063 |
| GeneID:7439 | 0.0003186807804837892 | GeneID:7430 | -0.0002556078056872143 |
| GeneID:10381 | 0.0003174085770658916 | GeneID:55793 | -0.0002558848103986634 |
| GeneID:55737 | 0.00031620944375184725 | GeneID:29928 | -0.0002562121802701867 |
| GeneID:10383 | 0.0003153799497778071 | GeneID:9929 | -0.00025666161799732144 |
| GeneID:10552 | 0.0003122251689802513 | GeneID:23408 | -0.00025747222936345603 |
| GeneID:10732 | 0.00031007641498102063 | GeneID:54504 | -0.00025965181798654087 |
| GeneID:23077 | 0.0003078562519840672 | GeneID:6674 | -0.0002601844105004172 |
| GeneID:25804 | 0.0003076865722904266 | GeneID:22911 | -0.0002632359652195597 |
| GeneID:6261 | 0.00030679586317539624 | GeneID:6533 | -0.00026349799376778244 |
| GeneID:404672 | 0.00030608988245862153 | GeneID:7862 | -0.00026625014118050836 |
| GeneID:23594 | 0.0003021986541538205 | GeneID:642 | -0.00026662265190429904 |
| GeneID:5971 | 0.00029964370273934154 | GeneID:79651 | -0.0002676439253825327 |
| GeneID:81848 | 0.00029588257669001684 | GeneID:9862 | -0.0002680283508349592 |
| GeneID:23348 | 0.0002945500638849357 | GeneID:26018 | -0.00027047788380515046 |
| GeneID:55651 | 0.0002934386491512662 | GeneID:6904 | -0.0002736807253864681 |
| GeneID:25940 | 0.0002907318435333681 | GeneID:112611 | -0.00027716501797302515 |
| GeneID:10574 | 0.0002845476716444732 | GeneID:79867 | -0.00027791256273626157 |
| GeneID:2201 | 0.00028438902519003345 | GeneID:4001 | -0.0002819600763266954 |
| GeneID:63895 | 0.0002838128604232874 | GeneID:8417 | -0.0002833042577312835 |
| GeneID:55666 | 0.0002819438923753184 | GeneID:4678 | -0.00028337110030636465 |
| GeneID:80271 | 0.0002818991331659287 | GeneID:23075 | -0.0002847791220921523 |
| GeneID:26499 | 0.0002797155758109364 | GeneID:27339 | -0.0002869081999755051 |
| GeneID:7248 | 0.00027835654026509216 | GeneID:54554 | -0.00028707619802396576 |
| GeneID:292 | 0.00027730360518614497 | GeneID:23011 | -0.0002873733538876851 |

| | | | |
|---|---|---|---|
| GeneID:25949 | 0.000277094023921693 | GeneID:5037 | -0.00028875808411212326 |
| GeneID:3185 | 0.00027663577545518144 | GeneID:5696 | -0.0002888639466219454 |
| GeneID:3028 | 0.0002758245191811366 | GeneID:51026 | -0.0002903371476901049 |
| GeneID:199 | 0.00027499707904491994 | GeneID:746 | -0.00029504257352229316 |
| GeneID:7726 | 0.0002746186666142056 | GeneID:6693 | -0.00029679994013166554 |
| GeneID:59343 | 0.0002724388964547083 | GeneID:529 | -0.00029723583942685227 |
| GeneID:1124 | 0.0002723803070153473 | GeneID:29104 | -0.0002981346741123123 |
| GeneID:81554 | 0.0002708230597962987 | GeneID:6993 | -0.00029819081847451027 |
| GeneID:4724 | 0.0002666732930872493 | GeneID:54020 | -0.00029838412984473415 |
| GeneID:55841 | 0.0002663926250959751 | GeneID:79053 | -0.00030115854376564513 |
| GeneID:2769 | 0.0002658547944375328 | GeneID:5092 | -0.0003054734205912938 |
| GeneID:10095 | 0.000264939410547679 | GeneID:5261 | -0.0003072767493442332 |
| GeneID:22933 | 0.00026460352244546946 | GeneID:80746 | -0.0003091011241485384 |
| GeneID:57214 | 0.00026348814658941696 | GeneID:8636 | -0.0003101706109962096 |
| GeneID:2783 | 0.0002617748084926127 | GeneID:126321 | -0.0003144495989148965 |
| GeneID:5698 | 0.00026107337450141665 | GeneID:23530 | -0.0003180070804040382 |
| GeneID:6443 | 0.0002598119608832808 | GeneID:256364 | -0.00031891705054651554 |
| GeneID:11180 | 0.00025957667675911974 | GeneID:10921 | -0.0003223289147302725 |
| GeneID:51650 | 0.0002593204003898797 | GeneID:716 | -0.000322608178093432 |
| GeneID:3265 | 0.0002592667936421793 | GeneID:51278 | -0.0003259721718142552 |
| GeneID:79017 | 0.00025884852084566087 | GeneID:902 | -0.00033134746579535797 |
| GeneID:648 | 0.000258685049081244 | GeneID:9778 | -0.0003313896104479341 |
| GeneID:445 | 0.00025666535832682096 | GeneID:2356 | -0.00033508541320056154 |
| GeneID:9404 | 0.00025395498832191934 | GeneID:10542 | -0.00033562939042893196 |
| GeneID:7041 | 0.0002523205544312143 | GeneID:2524 | -0.00033760608355564 |
| GeneID:116985 | 0.0002520070168574069 | GeneID:1021 | -0.0003378433632986821 |
| GeneID:26063 | 0.0002515512748850723 | GeneID:29761 | -0.00034363297259283214 |
| GeneID:6634 | 0.0002501185647212259 | GeneID:5469 | -0.0003474179997338746 |
| GeneID:7070 | 0.0002498839591129629 | GeneID:54665 | -0.0003526452723477166 |
| GeneID:25844 | 0.00024975960366947287 | GeneID:51703 | -0.0003645450562149516 |
| GeneID:79001 | 0.00024974034757776375 | GeneID:10939 | -0.0003700582783608668 |
| GeneID:2542 | 0.00024846847156446907 | GeneID:3915 | -0.0003750396714760705 |
| GeneID:4193 | 0.00024769030301595045 | GeneID:8446 | -0.00037538815318446575 |
| GeneID:1789 | 0.0002457226616798318 | GeneID:9044 | -0.0003767583503481191 |
| GeneID:1556 | 0.0002437985367657615 | GeneID:81566 | -0.0003878085662785802 |
| GeneID:51493 | 0.00024360876455965555 | GeneID:91369 | -0.0004094507881797229 |
| GeneID:49855 | 0.0002434010027060879 | GeneID:1759 | -0.0004395845189518063 |
| GeneID:8476 | 0.0002429233375216671 | GeneID:51646 | -0.00044154668812577795 |
| GeneID:8140 | 0.00024224934641965242 | GeneID:155066 | -0.000448837306106213 |

**Supplementary Table 3. GO enrichment analysis of biological processes for the prediction of kanamycin and paralysis.** Top 20 clusters with their representative enriched terms (one per cluster) created with Metascape. "Count" is the number of genes in the user-provided lists with membership in the given ontology term. "%" is the percentage of all of the user-provided genes that are found in the given ontology term (only input genes with at least one ontology term annotation are included in the calculation). "Log10(P)" is the p-value in log base 10. "Log10(q)" is the multi-test adjusted p-value in log base 10.

| GO | Description | Count | % | $Log_{10}(p)$ | $Log_{10}(q)$ |
|---|---|---|---|---|---|

| | | | | | |
|---|---|---|---|---|---|
| GO:0006974 | cellular response to DNA damage stimulus | 20 | 10.05 | -6.92 | -2.85 |
| GO:0007420 | brain development | 20 | 10.05 | -6.74 | -2.85 |
| GO:0097421 | liver regeneration | 5 | 2.51 | -5.91 | -2.20 |
| GO:0070542 | response to fatty acid | 6 | 3.02 | -5.42 | -1.84 |
| GO:0008283 | cell population proliferation | 16 | 8.04 | -4.75 | -1.47 |
| GO:0051301 | cell division | 13 | 6.53 | -4.43 | -1.24 |
| GO:1903047 | mitotic cell cycle process | 13 | 6.53 | -4.26 | -1.18 |
| GO:1901654 | response to ketone | 8 | 4.02 | -4.17 | -1.18 |
| GO:0030902 | hindbrain development | 7 | 3.52 | -4.17 | -1.18 |
| GO:0006869 | lipid transport | 10 | 5.03 | -4.15 | -1.18 |
| GO:0006739 | NADP metabolic process | 4 | 2.01 | -4.04 | -1.15 |
| GO:0009410 | response to xenobiotic stimulus | 11 | 5.53 | -3.94 | -1.08 |
| GO:0035272 | exocrine system development | 4 | 2.01 | -3.81 | -1.00 |
| GO:0055086 | nucleobase-containing small molecule metabolic process | 13 | 6.53 | -3.81 | -1.00 |
| GO:1901137 | carbohydrate derivative biosynthetic process | 13 | 6.53 | -3.79 | -1.00 |
| GO:0006914 | autophagy | 9 | 4.52 | -3.74 | -1.00 |
| GO:0009314 | response to radiation | 11 | 5.53 | -3.70 | -0.99 |
| GO:1903432 | regulation of TORC1 signaling | 4 | 2.01 | -3.55 | -0.95 |
| GO:0097305 | response to alcohol | 8 | 4.02 | -3.54 | -0.95 |
| GO:0007098 | centrosome cycle | 5 | 2.51 | -3.53 | -0.95 |

**Supplementary Table 4.** Genes of gene expression signature from kanamycin that were found to be highly positively or negatively attributed and their attribution score. The top 100 negatively and positively attributed genes are listed.

| Entrez Gene ID (positively attributed) | IG attribute score | Entrez Gene ID (negatively attributed) | IG attribute score |
|---|---|---|---|
| GeneID:1027 | 4,39E+07 | GeneID:6726 | -1,86E+10 |
| GeneID:7503 | 3,35E+10 | GeneID:7763 | -1,86E+10 |
| GeneID:29083 | 3,11E+10 | GeneID:1070 | -1,86E+10 |
| GeneID:23352 | 3,11E+10 | GeneID:4605 | -1,86E+10 |
| GeneID:10447 | 3,09E+10 | GeneID:3460 | -1,86E+10 |
| GeneID:8733 | 3,07E+10 | GeneID:26119 | -1,86E+10 |
| GeneID:1666 | 3,05E+10 | GeneID:137886 | -1,87E+09 |

| | | | |
|---|---|---|---|
| GeneID:3382 | 3,00E+10 | GeneID:7515 | -1,87E+10 |
| GeneID:7107 | 2,87E+10 | GeneID:6601 | -1,87E+10 |
| GeneID:7319 | 2,83E+09 | GeneID:6560 | -1,88E+10 |
| GeneID:11133 | 2,81E+10 | GeneID:55567 | -1,88E+10 |
| GeneID:7257 | 2,78E+10 | GeneID:55003 | -1,89E+10 |
| GeneID:112611 | 2,73E+09 | GeneID:5763 | -1,89E+10 |
| GeneID:4924 | 2,67E+10 | GeneID:157680 | -1,89E+10 |
| GeneID:633 | 2,65E+10 | GeneID:2235 | -1,90E+10 |
| GeneID:25805 | 2,60E+09 | GeneID:54386 | -1,91E+10 |
| GeneID:2958 | 2,58E+10 | GeneID:5891 | -1,91E+10 |
| GeneID:51290 | 2,55E+10 | GeneID:4967 | -1,92E+10 |
| GeneID:533 | 2,52E+09 | GeneID:2063 | -1,92E+10 |
| GeneID:8672 | 2,51E+10 | GeneID:5361 | -1,93E+09 |
| GeneID:8744 | 2,50E+09 | GeneID:6414 | -1,94E+10 |
| GeneID:10797 | 2,47E+10 | GeneID:883 | -1,95E+09 |
| GeneID:843 | 2,46E+10 | GeneID:5327 | -1,96E+10 |
| GeneID:2548 | 2,44E+10 | GeneID:51335 | -1,97E+10 |
| GeneID:5708 | 2,39E+09 | GeneID:1213 | -1,97E+10 |
| GeneID:4329 | 2,37E+09 | GeneID:51131 | -1,97E+10 |
| GeneID:201229 | 2,34E+10 | GeneID:10419 | -1,98E+10 |
| GeneID:10966 | 2,34E+09 | GeneID:1870 | -1,98E+10 |
| GeneID:63924 | 2,30E+10 | GeneID:51805 | -2,00E+09 |
| GeneID:3712 | 2,29E+10 | GeneID:10309 | -2,00E+09 |
| GeneID:11011 | 2,29E+10 | GeneID:54978 | -2,01E+09 |
| GeneID:1537 | 2,29E+09 | GeneID:3489 | -2,02E+10 |
| GeneID:5058 | 2,28E+09 | GeneID:63895 | -2,03E+10 |
| GeneID:8382 | 2,24E+07 | GeneID:1983 | -2,03E+09 |
| GeneID:8061 | 2,22E+09 | GeneID:8312 | -2,05E+10 |
| GeneID:5347 | 2,20E+09 | GeneID:348995 | -2,05E+09 |
| GeneID:55898 | 2,18E+10 | GeneID:29893 | -2,06E+08 |
| GeneID:54872 | 2,14E+09 | GeneID:51366 | -2,06E+09 |
| GeneID:7398 | 2,13E+09 | GeneID:10732 | -2,08E+10 |
| GeneID:22985 | 2,11E+09 | GeneID:55013 | -2,08E+10 |
| GeneID:28992 | 2,08E+10 | GeneID:25837 | -2,10E+09 |
| GeneID:5423 | 2,07E+10 | GeneID:144404 | -2,10E+10 |
| GeneID:978 | 2,04E+08 | GeneID:4666 | -2,10E+10 |
| GeneID:11215 | 2,04E+10 | GeneID:26151 | -2,11E+10 |
| GeneID:55329 | 2,04E+10 | GeneID:28978 | -2,12E+09 |
| GeneID:221061 | 2,02E+10 | GeneID:9040 | -2,13E+10 |
| GeneID:5649 | 2,02E+09 | GeneID:1374 | -2,14E+10 |
| GeneID:2539 | 2,01E+10 | GeneID:58488 | -2,14E+09 |
| GeneID:8925 | 1,98E+09 | GeneID:80254 | -2,14E+09 |
| GeneID:5226 | 1,98E+09 | GeneID:124222 | -2,15E+10 |
| GeneID:80758 | 1,98E+10 | GeneID:51195 | -2,19E+10 |
| GeneID:79084 | 1,97E+10 | GeneID:5355 | -2,21E+09 |
| GeneID:11060 | 1,96E+09 | GeneID:26001 | -2,21E+10 |
| GeneID:5727 | 1,93E+10 | GeneID:2203 | -2,26E+10 |
| GeneID:5111 | 1,91E+10 | GeneID:8428 | -2,26E+09 |
| GeneID:51026 | 1,91E+10 | GeneID:375056 | -2,27E+09 |
| GeneID:8624 | 1,91E+10 | GeneID:6284 | -2,33E+10 |
| GeneID:23231 | 1,90E+10 | GeneID:4983 | -2,34E+09 |

| | | | |
|---|---|---|---|
| GeneID:79006 | 1,90E+10 | GeneID:8506 | -2,34E+10 |
| GeneID:2218 | 1,90E+09 | GeneID:64759 | -2,35E+10 |
| GeneID:9181 | 1,90E+10 | GeneID:8514 | -2,35E+10 |
| GeneID:6867 | 1,89E+10 | GeneID:7534 | -2,36E+10 |
| GeneID:2564 | 1,89E+10 | GeneID:6286 | -2,37E+09 |
| GeneID:473 | 1,86E+10 | GeneID:1051 | -2,39E+09 |
| GeneID:10542 | 1,85E+10 | GeneID:55041 | -2,39E+09 |
| GeneID:27109 | 1,85E+10 | GeneID:10036 | -2,42E+09 |
| GeneID:51569 | 1,81E+10 | GeneID:653639 | -2,45E+10 |
| GeneID:57016 | 1,80E+10 | GeneID:7045 | -2,46E+10 |
| GeneID:10140 | 1,80E+10 | GeneID:1774 | -2,49E+10 |
| GeneID:65265 | 1,80E+10 | GeneID:6693 | -2,54E+10 |
| GeneID:5813 | 1,79E+10 | GeneID:4350 | -2,55E+10 |
| GeneID:55748 | 1,79E+09 | GeneID:51123 | -2,57E+10 |
| GeneID:57194 | 1,77E+10 | GeneID:7296 | -2,58E+10 |
| GeneID:9639 | 1,76E+09 | GeneID:10247 | -2,62E+10 |
| GeneID:2975 | 1,76E+08 | GeneID:9088 | -2,62E+10 |
| GeneID:4828 | 1,76E+10 | GeneID:5743 | -2,66E+10 |
| GeneID:4293 | 1,75E+10 | GeneID:28957 | -2,68E+08 |
| GeneID:64208 | 1,75E+10 | GeneID:400949 | -2,70E+10 |
| GeneID:6832 | 1,74E+09 | GeneID:6118 | -2,83E+10 |
| GeneID:9887 | 1,73E+10 | GeneID:57513 | -2,86E+10 |
| GeneID:29978 | 1,73E+10 | GeneID:10476 | -2,87E+10 |
| GeneID:10174 | 1,72E+09 | GeneID:54982 | -2,88E+09 |
| GeneID:83440 | 1,71E+10 | GeneID:7358 | -2,89E+10 |
| GeneID:4781 | 1,71E+10 | GeneID:4487 | -2,98E+10 |
| GeneID:1019 | 1,71E+10 | GeneID:51389 | -2,98E+10 |
| GeneID:51227 | 1,69E+10 | GeneID:2176 | -2,99E+10 |
| GeneID:10651 | 1,68E+10 | GeneID:5127 | -3,03E+10 |
| GeneID:79177 | 1,68E+10 | GeneID:10388 | -3,04E+09 |
| GeneID:9187 | 1,67E+10 | GeneID:5898 | -3,19E+09 |
| GeneID:6602 | 1,67E+10 | GeneID:9854 | -3,19E+10 |
| GeneID:5638 | 1,66E+10 | GeneID:80212 | -3,23E+09 |
| GeneID:64419 | 1,66E+10 | GeneID:8208 | -3,23E+10 |
| GeneID:9486 | 1,66E+10 | GeneID:8031 | -3,52E+10 |
| GeneID:6566 | 1,65E+10 | GeneID:10099 | -3,55E+10 |
| GeneID:9540 | 1,65E+10 | GeneID:79947 | -3,62E+10 |
| GeneID:26155 | 1,64E+10 | GeneID:3773 | -3,63E+10 |
| GeneID:55207 | 1,63E+10 | GeneID:1869 | -4,88E+09 |
| GeneID:4815 | 1,62E+10 | GeneID:5547 | -4,93E+09 |
| GeneID:84747 | 1,61E+10 | GeneID:6839 | -5,30E+09 |
| GeneID:80742 | 1,61E+10 | GeneID:2131 | -5,74E+09 |

**Supplementary Table 5.** Count of edge types in biomedical KG.

| Edge Type | Count |
|---|---|
| side-effect | 197 |

| | |
|---|---|
| functional association | 1761 |
| protein-condition | 5448 |
| physical interaction | 6087 |
| drug-protein | 12451 |
| indication | 12848 |
| genetic protein-condition association | 159202 |

**Supplementary Table 6.** Edge type count in biomedical KG according to source database.

| **Source Database** | **Count** |
|---|---|
| PHEWAS | 159202 |
| BIOGRID | 102447 |
| KEGG | 63356 |
| PATHWAYCOMMONS | 32928 |
| INTACT | 23055 |
| DRUGBANK | 10072 |
| CLINICALTRIALS | 7626 |
| REACTOME | 6379 |
| DISGENET | 5448 |
| OPENTARGETS | 5222 |
| IUPHAR | 2379 |
| NEUROMMSIG | 1761 |
| SIDER | 135 |
| OFFSIDES | 62 |

**Supplementary Table 7.** Hyperparameters selected by the optuna hyperparameter optimization for the MultiGML-variants (-RGCN and -RGAT) and the respective ranges for each hyperparameter.

| **Hyperparameter Name** | **Range** | **Best Value RGCN** | **Best Value RGAT** |
|---|---|---|---|
| Input dimension | [10, 500; 10] | 390 | 8 |
| Hidden dimension | [10, 300; 10] | 280 | 200 |
| Output dimension | [10, 100; 10] | 60 | 24 |

| | | | |
|---|---|---|---|
| Number of hidden layers | [1 : 7] | 3 | 2 |
| Dropout embedding layer | [0.0, 0.4; 0.1] | 0.4 | 0.2 |
| Dropout graph neural network | [0.0, 0.5; 0.1] | 0.0 | 0.1 |
| Learning rate (optimizer) | [10e-5 : 10e-2] | 0.00028 | 0.00024 |
| Weight decay (optimizer) | [1e-4, 1e-5, 1e-6] | 1e-06 | 1e-05 |
| Epsilon | [1e-8 : 1e-6] | 5.85e-07 | 6.12e-07 |
| Batch size | [100, 800; 100] | 100 | 300 |
| Drug modalities | all possible combinations of [gene expression signature, cytological profiling signature, molecular fingerprint] | drug_fc, drug_fcfp | drug_fcfp |
| Protein modalities | all possible combinations of [gene ontology fingerprint, protein sequence embedding] | protein_go | protein_go |
| Number of attention heads | [1, 2, 3, 4] | - | 2 |
| Dropout graph attention network | [0.0, 0.5; 0.1] | - | 0.0 |
